# High heritabilities for antibiotic usage show potential to breed for disease resistance in finishing pigs

**DOI:** 10.1101/2021.06.17.448802

**Authors:** Wim Gorssen, Dominiek Maes, Roel Meyermans, Jürgen Depuydt, Steven Janssens, Nadine Buys

## Abstract

The use of antimicrobials in animal production is under public debate, mainly due to the risk of transfer of resistance to pathogenic bacteria in humans. Therefore, measures have been taken during the last decades to reduce antibiotic usage in animals, for instance by national monitoring programmes and by improving animal health management. Although some initiatives exist in molecular genetic selection, quantitative genetic selection of animals towards decreased antibiotic usage is an underexplored area to reduce antibiotic usage. However, this strategy could yield cumulative effects.

In this study we derived new phenotypes from on-farm parenteral antibiotic records from commercially grown crossbred finishing pigs used in the progeny test of Piétrain terminal sires to investigate the heritability of antibiotics usage. Parenteral antibiotic records, production parameters and pedigree records of 2238 fullsib pens from two experimental farms in Belgium between 2014 and 2020 were analysed. Heritability estimates were moderate (18-44%) for phenotypes derived from all antibiotic treatments, and low (1-15%) for phenotypes derived from treatments against respiratory diseases only. Moreover, genetic correlations between these new phenotypes and mortality were low to moderate (0.08-0.60) and no strong adverse genetic correlations with production traits were found.

The high heritabilities and favourable genetic correlations suggest these new phenotypes derived from on-farm antibiotics records to be promising for inclusion in future pig breeding programs to breed for a decrease in antibiotics usage.

## 1. Introduction

Antimicrobials are commonly used in animal production systems, not only for treatment, but also for metaphylaxis, prophylaxis and in some countries still for improvement of feed efficiency and growth (1,2). However, antimicrobial usage is under public debate, mainly because of the risk of selecting resistant bacteria in animals and the transfer of resistance to pathogenic bacteria in humans (1,3). Therefore, legislation has been developed in many countries to restrict the use of antimicrobials in animals. In the European Union (EU), the use of antibiotics for growth promotion has been banned since 2006 (Regulation (EC) No 1831/2003 of the European Parliament and of the Council on additives for use in animal nutrition). Furthermore, the EU commission goal is to reduce antimicrobials in livestock and aquaculture by 50% between 2020 and 2030, according to the ‘From Farm to Fork’ strategy of the European Green Deal (4). These incentives have already led to a significant decrease in antibiotic usage in the EU and studies have shown that this is feasible without production losses (5,6). Between 2011 and 2017, the sales of antibiotics for animals in the EU were reduced by more than 32% (7). This reduction was mainly achieved by three strategies: (i) implementing monitoring systems for quantifying antimicrobial usage, (ii) allowing benchmarking by restricting the use of critically important antibiotics such as fluoroquinolones and third and fourth generation cephalosporins (8) and (iii) encouraging other control and preventive measures against infections improving animal health management (3,5). The latter measures mainly relate to optimising management and biosecurity, housing conditions, nutrition and by implementing vaccination. However, each of these measures also have limitations and/or cannot always be implemented easily in farms (9,10).

Reducing antibiotic usage in animals by genetic selection has been underexplored so far (11). Breeding animals for a decreased antimicrobial usage could improve general disease resistance and could yield cumulative and permanent gains (12). Breeding for higher disease resistance also fits very well within the concept “Raised Without Antibiotics (RWA)”, a certification mark that is increasingly promoted and implemented in different countries. The RWA protocol implies that no antibiotics of any kind are used in raising the animals. Previous studies in pigs mainly focused on breeding for resistance against specific diseases (13–15) or for favourable immunological trait levels (16,17). Breeding pigs for resistance towards specific diseases may however lead to increased susceptibility for other diseases (13,15). Studies focusing on immunological traits, such as viral load or levels of cytokines and antibodies, reported moderate to high heritabilities and a clear link with animal health (16,18). Nonetheless, phenotyping immunological traits is costly and challenging to incorporate in a breeding program (19). Furthermore, simple binary (*e.g*. treated vs non-treated) or ordinal (*e.g*. none, mild or severely diseased) recordings of pig health status might be insufficient for breeding programs (19). For example, very low heritabilities (4-6%) were reported by Guy et al. (11) for binary health traits (medicated vs non-medicated) in pigs. Therefore, new and quantitative phenotypes might be better suited to breed animals towards disease resistance. Henryon et al. (20) reported heritability estimates of 10-19% for time until first treatment on a logarithmic scale for different disease categories, whereas Putz et al. (21) found a heritability of 13-29% for number of antibiotic treatments at individual pig level. However, these studies using field data only focused on the number of treatments pigs received.

In this study, we investigate the value of field data on antibiotic usage in finishing pigs on a pen level as a new, easy to measure phenotype in pig breeding. Genetic parameters such as heritability and genetic correlations were estimated for antimicrobial usage in finishing pigs.

## 2. Materials and Methods

### Description of study population

Data were collected on crossbred progeny of Piétrain sires and hybrid sows (Large White/Landrace crosses, Danbred and Topigs 20) kept in two experimental fattening units (farm A and B) of the breeding organization “Vlaamse Piétrain Fokkerij” (VPF, Belgium). After quality control, data were available on 2238 full-sib pens with known pedigree (14742 finishing pigs; Table 1 and 2). Pigs entered the experimental farms at a median age of 68 days (24.0±4.1kg) and were slaughtered at a median age of 191 days (114.0±9.3kg) (Table 2). Piglets were vaccinated against *Mycoplasma hyopneumoniae* before weaning and received anthelminthic medication upon arrival at the experimental farm. Pigs were kept in commercial conditions representative for Belgian pig production (*e.g*. semi slatted floor, ad libitum pellet feeding, drinking nipple) but were kept in pens of only 6 to 7 full-sib pigs. Farm A had 18 compartments, each containing 8 pens (4.7 m^2^ per pen) of usually 6 pigs. Pens in farm A were of mixed sex with females and immunocastrated males to prevent boar taint (Improvac®, Zoetis). Farm B had 7 compartments, each with 16 pens (5.4 m^2^) of usually 7 pigs. Pens in farm B were of mixed sex composed of females and surgically castrated males (barrows). Only pens starting with 5 to 8 full-sib pigs were retained for further analyses.

**Table 1.** Descriptive statistics of traits at pen level for antibiotics usage parameters and other production traits (final data, N= 2238 pens).

| Trait | mean | sd | median | min | max |
| --- | --- | --- | --- | --- | --- |
| Number pigs per pen at start | 6.6 | 0.6 | 7.0 | 5.0 | 8.0 |
| Proportion of female pigs pen | 0.5 | 0.19 | 0.5 | 0.0 | 1.0 |
| UD<sub>Pen-all</sub> (mg/kg) | 1.07 | 2.79 | 0.00 | 0.00 | 41.60 |
| UD<sub>Pen-respiratory</sub> (mg/kg) | 0.57 | 2.11 | 0.00 | 0.00 | 41.60 |
| TI<sub>ADD-all</sub> | 0.11 | 0.31 | 0.00 | 0.00 | 4.32 |
| TI<sub>ADD-respiratory</sub> | 0.07 | 0.24 | 0.00 | 0.00 | 4.32 |
| TI<sub>UDD-all</sub> | 0.15 | 0.39 | 0.00 | 0.00 | 4.00 |
| TI<sub>UDD-respiratory</sub> | 0.09 | 0.30 | 0.00 | 0.00 | 3.84 |
| N<sub>treatment-all</sub> (days) | 0.7 | 1.5 | 0.0 | 0.0 | 19.0 |
| N<sub>treatment-respiratory</sub> (days) | 0.4 | 1.0 | 0.0 | 0.0 | 12.0 |
| AB<sub>ml-all</sub> (ml) | 3.4 | 8.4 | 0.0 | 0.0 | 88.0 |
| AB<sub>ml-respiratory</sub> (ml) | 2.1 | 6.9 | 0.0 | 0.0 | 88.0 |
| AB<sub>mg-all</sub> (mg) | 425.9 | 1121.5 | 0.0 | 0.0 | 12800.0 |
| AB<sub>mg-respiratory</sub> (mg) | 256.8 | 911.8 | 0.0 | 0.0 | 12800.0 |
| Mortality per pen (counts) | 0.08 | 0.31 | 0.00 | 0.00 | 4.00 |
| **FI (kg/day)** | 1.81 | 0.16 | 1.81 | 1.04 | 2.85 |
| **FCR** | 2.40 | 0.17 | 2.38 | 1.85 | 3.79 |

Abbreviations: UD: Used antibiotic dose; TI: Treatment incidence; N<sub>treatment</sub>: Treatment duration in days. AB<sub>mg</sub>:
Administered amount of active compound of antibiotics in mg; AB<sub>ml</sub>: Administered amount of product in ml; FI:
Feed intake; FCR: Feed conversion ratio. The suffix “–all” refers to the dataset with all parenteral antibiotic
treatments, whereas the suffix “–respiratory” refers to the dataset with only treatments for respiratory conditions.

**Table 2.** Descriptive statistics of production traits and covariables for analyses at individual level (final data, N=14742 pigs).

| Trait | mean | sd | median | min | max |
| --- | --- | --- | --- | --- | --- |
| Age at start (days) | 69.5 | 6.0 | 68.0 | 59.0 | 113.0 |
| Weight start (kg) | 23.9 | 4.1 | 24.0 | 12.0 | 51.0 |
| Age at slaughter (days) | 189.5 | 14.7 | 191.0 | 68.0 | 242.0 |
| Weight at slaughter/death (kg) | 114.0 | 9.3 | 114.0 | 30.0 | 147.1 |
| Carcass Weight (kg) | 90.4 | 6.6 | 90.2 | 44.9 | 120.1 |
| Duration test (days) | 120.0 | 14.8 | 120.0 | 0.0 | 168.0 |
| ADG (kg/day) | 0.759 | 0.093 | 0.758 | 0.341 | 1.149 |
| Meat% (%) | 62.9 | 2.7 | 63.1 | 45.4 | 70.2 |

Abbreviations: ADG: Average daily gain; Meat%: Lean meat percentage of the carcass.

### Data collection of antimicrobial usage

Antibiotic usage was registered at pen level only between February 2014 and February 2017 at farm A and between February 2014 and September 2020 at farm B. The choice of antimicrobial and the administration route were decided by animal caretakers upon evaluation of the pig health status and according to the guidelines of the herd veterinarian. Antimicrobials were only administered for treatment or metaphylaxis, not for prophylaxis. The following parameters were recorded: date of treatment, pen identification, reason of treatment, product name, number of treated pigs, administered dosage (ml per pig) and treatment duration (days). Other medication such as vaccines, anthelmintic products and anti-inflammatory drugs were not considered in this study.

### Data collection performance parameters

Aside from antimicrobial usage, also feed intake (FI) during the testing period was recorded at the pen level. Average daily gain (ADG), mortality during the finishing period and carcass quality traits (meat%) were collected at the individual level. Pigs that died before slaughter were individually weighed so that all weight data could be adjusted and retained in the analysis.

### Antibiotic usage parameters

Antimicrobial usage data were summarised per pen. In farm A, 104 pens were group medicated at the compartment level with an oral antibiotic (Soludox©, Eurovet Animal Health BV) on a total of 779 pens with at least one medication event (13.5%). These pens were excluded due to the different route of administration (oral vs injectable) of the antibiotic.

Two datasets were created: one dataset with all antibiotic treatments (AB_all_) and another set with antibiotic treatment for respiratory problems only (AB_respiratory_). The AB_respiratory_ dataset was created because the majority of antibiotics were used to treat respiratory diseases (70.5% of all treatments). In AB_respiratory_, treatments with fluoroquinolones (1 pen) and macrolides (7 pens) were rare and therefore these treatments were removed from the dataset. An overview of the antibiotics reported in this study is shown in Additional file 1 (Table S1).

The treatment length (N_treatment_) in a pen was calculated by summing the treatment duration in days from all treatments of that pen. Furthermore, the total antimicrobial usage in mg active compound (AB_mg_) was calculated as follows: first, the total administered volume of an antibiotic (AB_ml_) was calculated for each treatment:

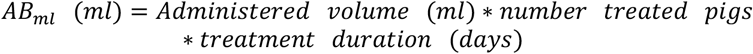

Second, AB_mg_ was computed using the concentration (mg/ml) of the active compound (Additional file 1 Table S1):

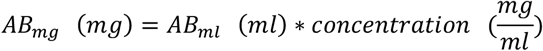

For trimethoprim-sulfonamides, the concentration of the minor substance trimethoprim was used (40 mg/ml), as suggested by Timmerman et al. (22).

Third, the used antibiotic dose (UD_Pen_; mg/kg) was calculated as mg of active compound administered per kg of live weight of pig in the pen at the time of administration (based on UDD from Timmerman et al. (22))

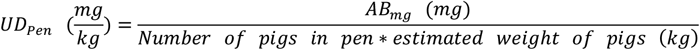

The weight at treatment was estimated based on the age of pigs using a Gompertz growth curve function:

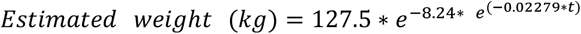

where *t* is the time at treatment (age in days). The growth curve parameters were estimated using the available weight data at start and at death or slaughter (Table 2).

The used daily dose per pig (UDD_pig_), the administered dose of a drug per kg pig per day, and the animal daily dose per pig (ADD_pig_) (23), the nationally recommended dose of a drug per kg pig per day, were calculated following Timmerman et al. (22) and Callens et al. (24). These parameters were used to assess whether dosing had been done according to label directions. Antibiotic administrations where 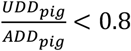 were considered as underdosed, whereas administrations were considered over-dosed when 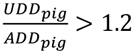 (22,24). Estimates of ADD_pig_ were based on dose recommendations by the Belgian expertise center on antimicrobial use in animals (AMCRA; 25). Treatment incidences (TI_ADD_ and TI_UDD_) were calculated at the pen level for each parenterally administered treatment (2):

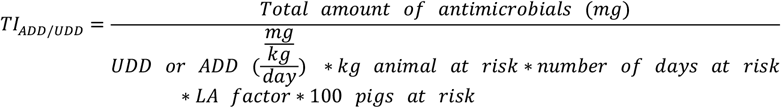

where the total amount of antimicrobials administered equals AB_mg_; UDD or ADD is as specified before; LA is a correction factor for long acting antibiotics (25); kg animal at risk is the estimated total weight of pigs in a pen; and number of days at risk was the pens’ test period (median 123 days). The TI_ADD_ and TI_UDD_ can be interpreted as the percentage of days in a period ‘at risk’ a pig is treated with one reference dose (ADDpig or UDD_pig_).

Finally, AB_ml_, AB_mg_, N_treatment,_ UD_pen_, TI_ADD_ and TI_UDD_ were summed per pen over all treatments (if the specific pen had multiple treatments) to obtain a dataset with one observation per pen, including all above mentioned traits. Pens where no antibiotics were used received a value of ‘0’. Traits for the full dataset AB_all_ received the suffix ‘-all’ (*e.g*. UD_pen-all_), whereas traits in the AB_respiratory_ dataset had a suffix ‘-respiratory’ (*e.g*. UDpen-respiratory).

### Data editing of performance traits

Average daily gain (ADG) was calculated individually as the difference between weight at slaughter/death and start of test (kg), divided by duration of test (days). The percentage of lean meat (meat%) was estimated using a fully automated classification system (AutoFom III(tm)) at two slaughterhouses from the Belgian Pork Group. Mortality counts were summed at the pen level. Feed conversion ratio (FCR) was calculated as the mean FI (kg/day) divided by the mean ADG (kg/day) per pen. Details of trait distributions are shown in Table 1 and Table 2.

The final dataset after data cleaning included 2238 pens: 14742 finishing pigs from 699 Piétrain sires and 1148 crossbred dams. Pedigree comprised 24081 animals, where the median pedigree depth of Piétrain sires was 11 generations (min 1; max 17) and 5 (min 0; max 11) for crossbred dams.

### Modelling

Heritabilities (h^2^) were estimated using single-trait genetic animal models with average information REML, implemented in *airemlf90* and invoked with the R-package *breedR* (26,27). For traits measured on the individual level (ADG and Meat%), each animal was analysed with corresponding pedigree. For traits measured on the pen level, full-sib pens were integrated in one record and treated as a single animal with corresponding pedigree. Genetic correlations (r_g_) between traits related to antibiotic usage (UD_Pen-all_, N_treatment-all_, UD_Pen-respiratory_ and N_treatment-respiratory_) were estimated using bivariate animal models (*airemlf90*). The h^2^ was estimated as the proportion of additive genetic variance divided by total variance, whereas c^2^ was estimated as the proportion of variance explained by random environmental effects (c), divided by total variance. Standard errors of h^2^, c^2^ and r_g_ were estimated using the *se_covar_function* option in *airemlf90*.

Animal models were of the form:

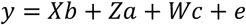

Where *y* is the vector with phenotypes for the studied trait(s); *b* is the vector containing the fixed effects (experimental farm, 2 levels; line of the sow, 2 levels; parity of dam, 10 levels) and covariates (number of pigs in pen; sex (individual level) or proportion of females in pen; (mean) duration of testing period); *a* is the vector of additive genetic effects (8520 levels); *c* is the vector of random environmental effects (179 levels); *e* is the vector of residual effects; *X, Z* and *W* are incidence matrices for respectively fixed effects, random animal effects and random permanent environmental effects. The random environmental effect *c* is a combination of farm, compartment and month of start of testing period (*e.g*. FarmA_Compartment1_2014_02) and was required to contain the progeny of at least 2 sires. The median number of pens in a common environmental effect was 8 and common environmental effects with less than 4 pens were merged in a remainder group (8 pens). Parity of sows ranged from 1 to 9 and parities above 8 were merged in a remainder group ‘9’.

### Cross-validation

The predictive ability of models for UD_Pen-all_, N_treatment-all_, UD_Pen-respiratory_ and N_treatment-respiratory_ was assessed using five-fold cross-validation with random masking of 20% of the data. Furthermore, 10 replications were made to avoid random sampling effects (28), so in total, 50 runs per trait were performed, allowing to also assess stability of h^2^ and c^2^ estimates.

Cross-validation was performed as follows: a univariate animal model (as specified before) was fitted on the complete datasets. From these results, observed phenotypes were adjusted for fixed and non-genetic random effects: 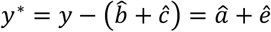. Predictive abilities were estimated as the Pearson correlation between breeding values of a validation dataset (with masked phenotypes) and the adjusted phenotypes (y*): *Predictive ability* = *r*(*EBV*_*masked*_, *y*^∗^). Two masking strategies were applied. First, one out of five pens was randomly masked within a sire family, that is within the pens of a sire. Second, one out of five pens was randomly masked across families, so all progeny from one out of five sires was masked. If there were less than five observations for a given sire, masking was performed using a probability of 20% for each observation, with maximum one masked observation per sire. The within-family strategy allows to determine predictive ability of breeding values mainly from close – half-sib – relationships, whereas across-family masking allows to determine predictive ability of more distant relationships, for example at the grandparent-level (28).

## 3. Results

### 3.1. Descriptive statistics

In total, 626 pens had a minimum of one pig treated, at least once, with antibiotics (28.0% of AB_all_ dataset). Most of the 1538 antibiotic treatments were from the class of lincosamides (44.5%), cephalosporins (18.4%), aminopenicillins (18.3%) and florfenicol (11.4%). The majority of treatments were applied for respiratory diseases (70.5%). Other indications included locomotion problems (8.7%), diarrhoea (5.6%), severe weight loss (4.5%), oedema disease (3.6%), ear infections (2.9%) and others (4.2%). The number of treatments per month is given in Figure 1.

**Figure 1.**
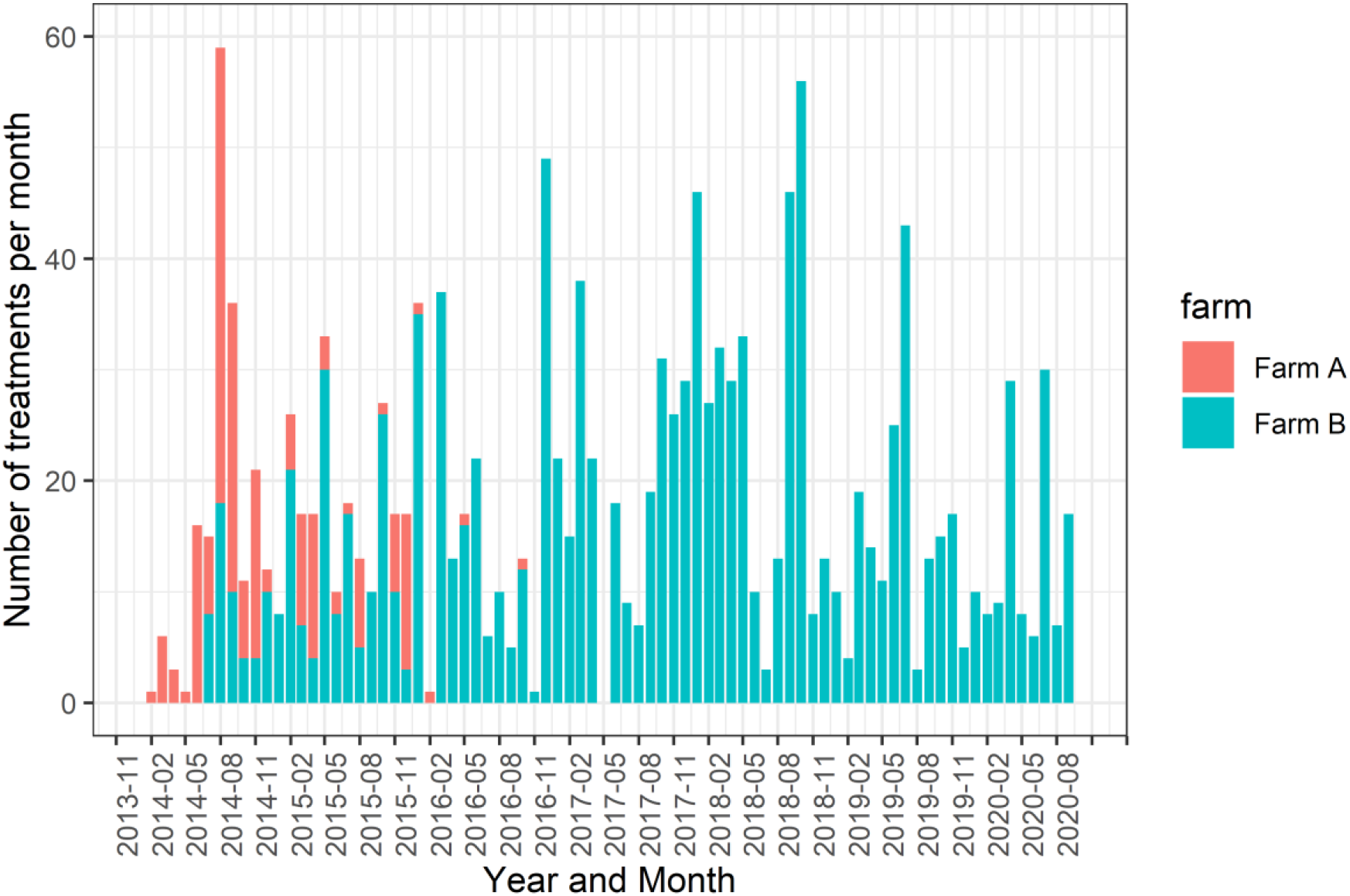
Number of parenteral antibiotic treatments per month for experimental farms A and B.

Descriptive statistics of the studied traits at a pen level are shown in Table 1, whereas descriptive statistics of traits studied at the individual level are shown in Table 2. Traits related to antibiotic usage (UD_pen_, N_treat-_ ment, AB_ml_, AB_mg_, TI_ADD_ and TI_UDD_) and mortality were zero-bound and showed a heavy right-tailed distribution. Other traits were approximately normally distributed based on visual inspection of trait distributions.

On average for the AB_all_ dataset, pens were treated 0.7 times with 3.6 ml of antibiotics and 445.1 mg of active compound (Table 1). The mean UD_pen-all_ was 1.27 mg of active compound administered per kg pig in the pen which is equivalent to a single administration of 127 mg of active compound to a fattening pig of 100 kg. The UDD/ADD ratio showed that 33.5% of treatments were correctly dosed (0.8 < UDD/ADD < 1.2), whereas 49.4% of treatments were underdosed and 17.1% were overdosed. Treatment incidences (TI_ADD-All_, TI_ADD-respiratory_, TI_UDD-All_, TI_UDD-respiratory_) ranged between 0.08 and 0.17, indicating that an average pig was treated with one reference dose of antibiotics for 0.08-0.17% of the period at risk (119 days median).

### 3.2. Genetic parameters

Estimates of h^2^, c^2^ and r_g_ are shown in Table 3. Heritabilities of traits related to antibiotic consumption were moderate to high for the full dataset (18-44%) whereas they were low to moderate for the respiratory problems dataset (1-15%). However, for both datasets, AB_mg_, TI_ADD_ and UD_pen_ were most heritable and significantly >0 based on a 95% confidence interval (mean ± 1.96*se), which is not explicitly given. In general, c^2^ was moderate to high for antibiotic usage (12-38%) and estimates were larger in the respiratory problems dataset. UD_pen_ and N_treatment_ across datasets were highly genetically correlated (r_g_=0.65 to 0.95). Genetic correlations with production traits were low with high standard errors, but the genetic correlation with mortality was consistently positive, ranging from 0.08 to 0.60.

**Table 3.**
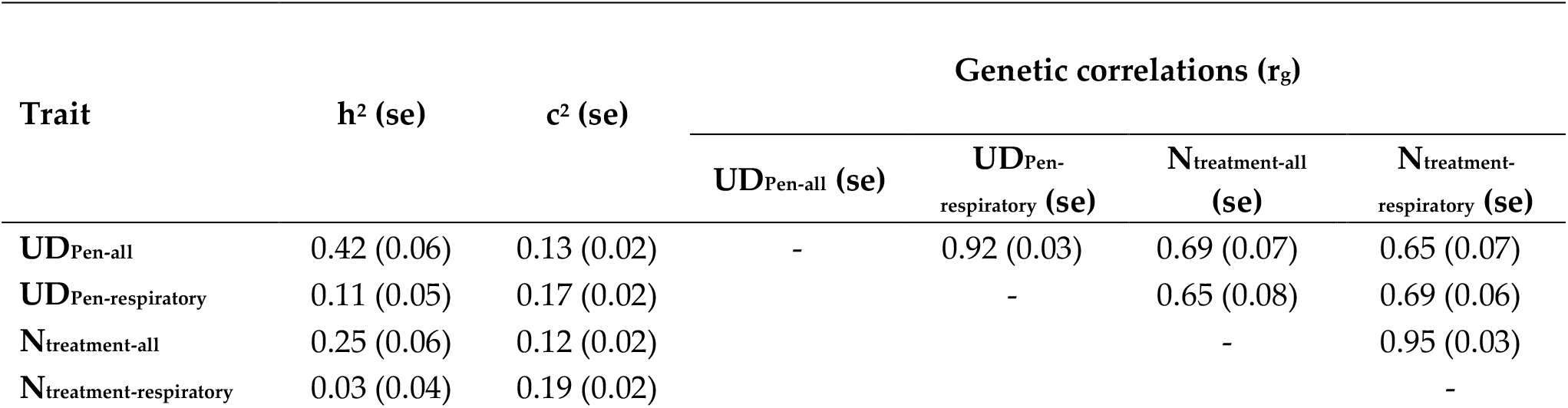
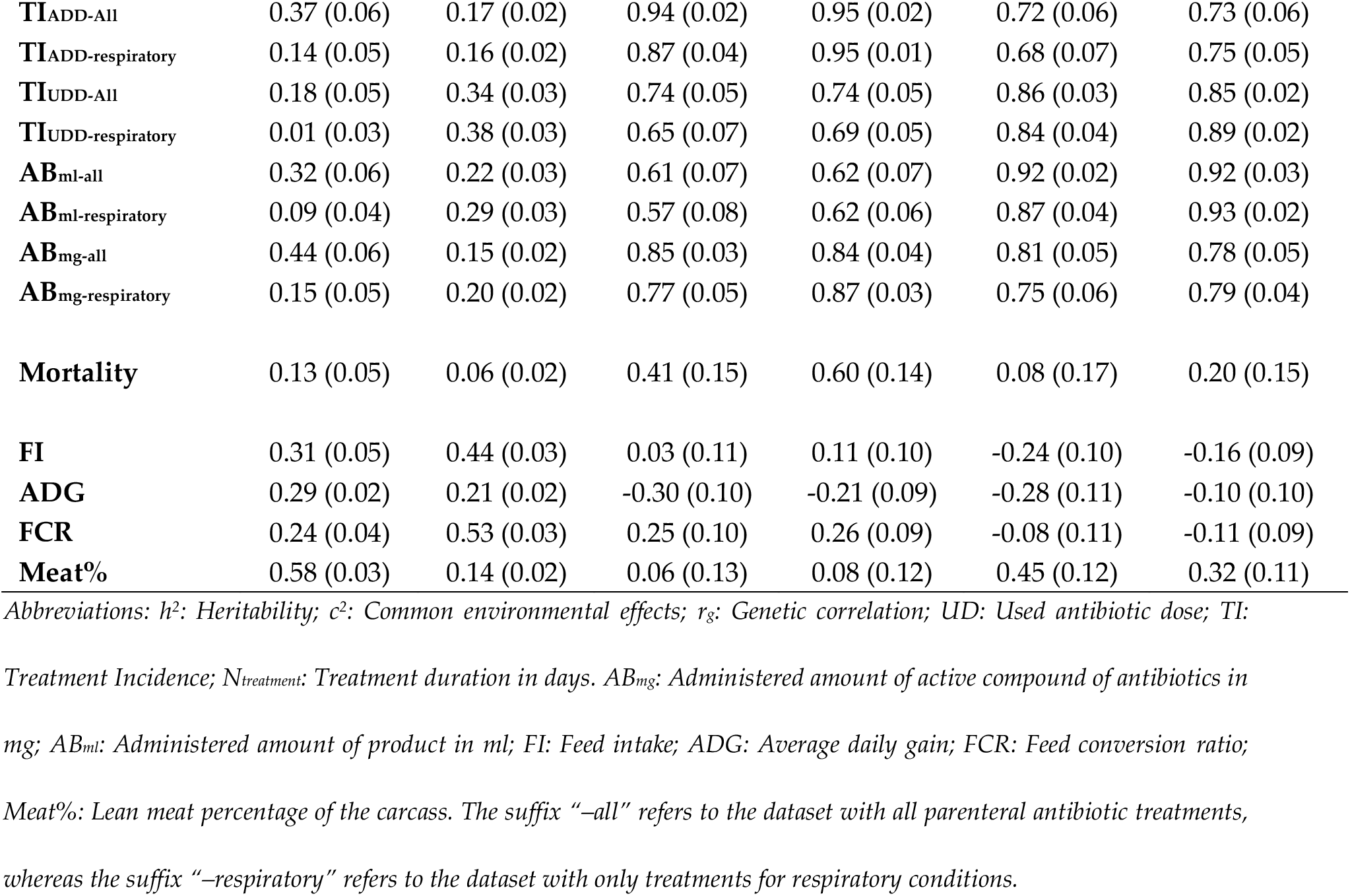
Genetic parameters for the antibiotic usage traits (pen level) and production traits (individual/pen level). Heritabilities (h^2^), common environmental effects (c^2^) and genetic correlations (r_g_; bivariate) were calculated.

### 3.3. Predictive abilities

Results of cross-validation analyses for UD_pen_ and N_treatment_ are shown in Table 4. Predictive abilities (given as Pearson correlations) were significantly higher than zero based on a 95% confidence interval, except for the AB_respiratory_ dataset using the across-family sampling strategy. Within-family sampling yielded – as expected due to the exploitation of close relationships – higher predictive abilities with lower standard errors. Predictive ability of UD_Pen_ was higher than for N_treatment_. Heritabilities obtained in cross-validations were similar to the full dataset.

**Table 4.**
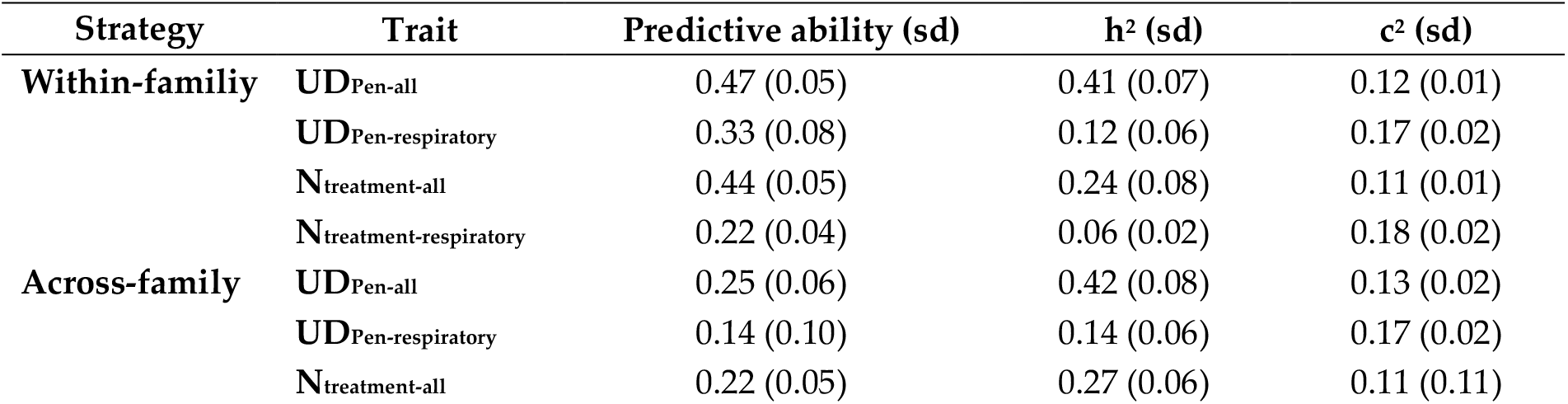
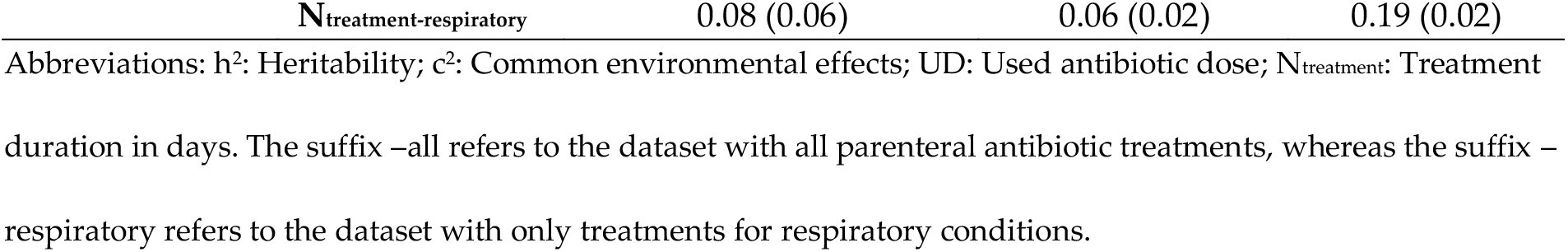
Cross-validation results for within-family and across-family cross-validation. Estimates of predictive abilities (correlation of EBVs from validation dataset with corrected phenotypes), heritabilities (h^2^) and common environmental effects (c^2^) are given.

## 4. Discussion

This study is the first to report genetic parameters of antibiotic usage derived from medication records in finishing pigs. New phenotypes for antibiotic usage at a pen level in finishing pigs were derived from on-farm medication records and were found to be moderately heritable and with a favourable (low to moderate) genetic correlation with mortality.

### Antibiotic usage

Six different phenotypes derived from medication records at pen level were investigated: AB_ml_, AB_mg_, N_treatment_, TI_ADD_, TI_UDD_ and UD_Pen_. Data were summed over different antibiotics at a pen level with an equal weight to the administered mg of active compound, although potency (strength) of active compound can differ among products (29). However, accounting for differences in potency via correction factors is delicate, also because this is partly corrected for with differences in recommended doses of a drug per kg pig per day. Therefore, we analysed traits with an equal weight on the administered mg of active compound.

When considering all treatments, 28.0% of all pens were treated at least once, and 19.7% of all pens were treated for respiratory problems. The UDD_pig_/ADD_pig_ ratios showed that most treatments were underdosed (61.9%), while 12.6% were overdosed and 25.5% correctly dosed (Figure 2). This is in contrast with previous studies in Belgian and Austrian pig herds, where overdosing was found to be more frequent for parenteral treatments (24,30). The dosage used by the farmer in the latter studies was based on visual assessment of the weight of the pigs, whereas in this study, weights were estimated based on age at treatment and using an estimated growth curve using all available weights. Possibly, treated pigs weighed on average less than their peers, increasing the UDD/ADD ratio and causing an underestimation of the correctness of dosing. Moreover, in this study only treatments in finishing pigs were investigated, whereas the studies referred to included also data on suckling and weaned piglets.

**Figure 2.**
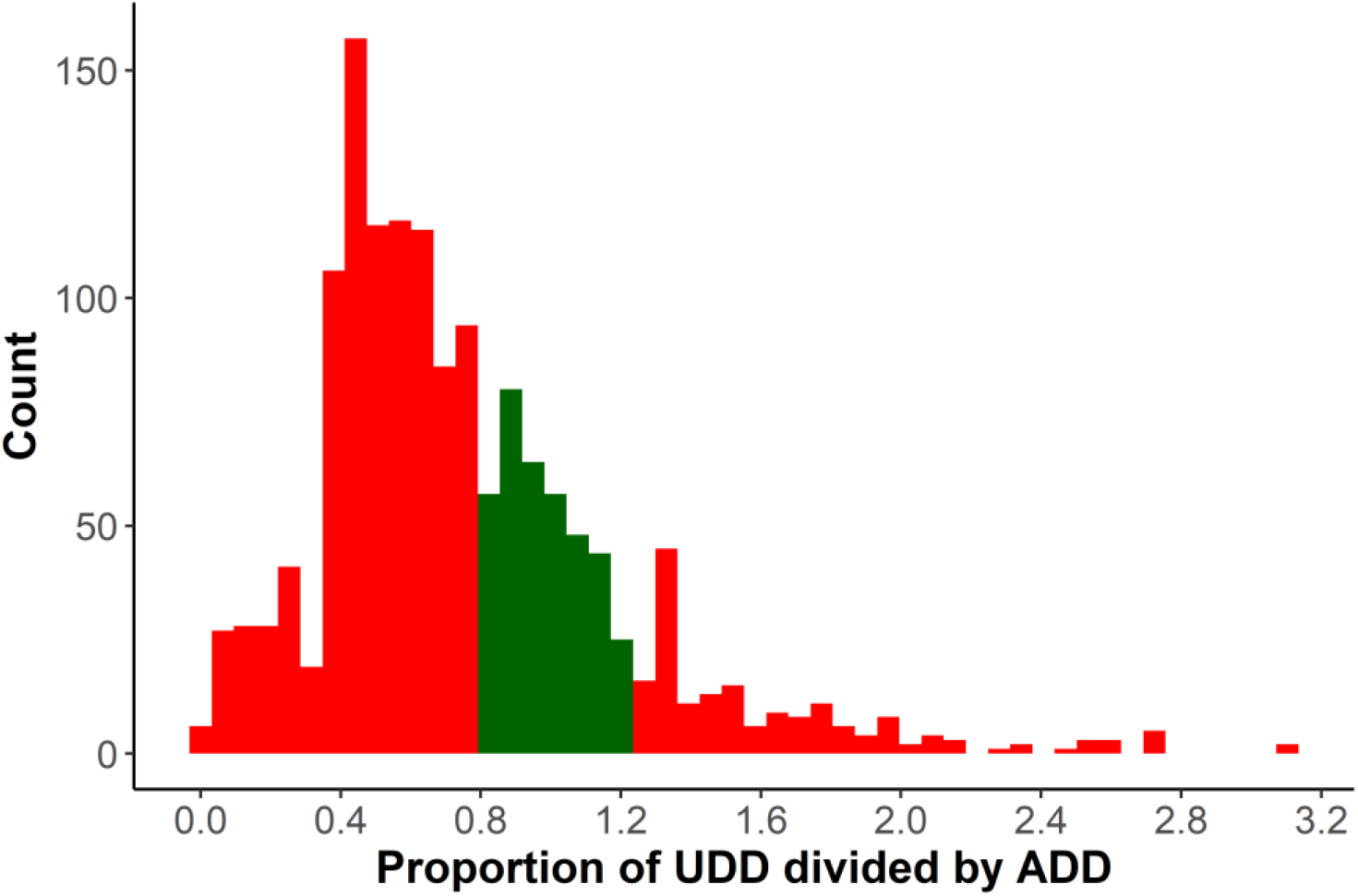
Histogram of UDD_pig_/ADD_pig_ ratio. Correctly dosed treatments are coloured green, whereas underdosed (<0.8) and overdosed (>1.2) treatments are coloured in red. These thresholds were set according to Callens et al. (2012). Abbreviations: UDD_pig_: Used daily dose per pig; ADD_pig_: Average daily dose per pig.

Treatment incidences (TI_ADD_ and TI_UDD_) were low, ranging from 0.09 to 0.15 units, meaning that pigs were treated with one reference dose of antibiotics for 0.09-0.15% during the period at risk (fattening period). Sjölund et al. (31) found median TI values of 0.00, 0.28, 0.83 and 2.09 for French, Swedish, Belgian and German fattening pig herds, respectively, and in a study on 9 different countries (partly same data) Sarrazin et al. (2) calculated a median TI of 1.2 for finishing pigs. Our estimates of TI_ADD_ and TI_UDD_ are thus rather low, especially compared to the median Belgian value of 0.83.

Figure 1 shows that parenteral treatments occurred relatively regular in time over farms, indicating that pigs frequently faced health issues and were challenged. The presence of an external challenge is critical to express genetic variability in antibiotic usage and disease resistance (15,20). In order to take spatiotemporal differences in infection pressure into account, the specific batch of pigs within a compartment (based on start date of testing) was included as a random environmental effect in genetic modelling. Indeed, it can be assumed that infection pressure will be relatively consistent within such a group, because of the close proximity of pigs in a confined space over a prolonged period (median 123 days, Table 2) (15).

### Genetic analysis of antibiotic usage

Most studies on genetic selection for health traits in various livestock species focused on binary (*e.g*. treated vs non treated) and/or ordinal traits (*e.g*. none, mild or severe symptoms), assuming that a higher health status would decrease antimicrobial usage and mortality (11,32,33). Furthermore, Henryon et al. (20) used time until first treatment (days) in pigs as a proxy for disease resistance, while Putz et al. (21) investigated the number of treatments in pigs. In our study, alternative phenotypes were derived based on the actual amount of parenterally administered antibiotics at a pen level. These new phenotypes are zero-bound quantitative traits which allow to quantify antibiotic treatments more in detail and, possibly, disease status. A more continuous phenotype has advantages over categorical data and we believe that our approach could refine genetic analyses and provide more accurate estimates.

Heritability (h^2^) estimates for traits related to antibiotic usage were moderate to high in the dataset containing all parenteral antibiotic treatments (18-44%) and low to moderate in the respiratory problems dataset (1-15%) (Table 3). The h^2^ was highest for AB_mg_, TI_ADD_ and UD_Pen_, indicating these parameters offer most potential to reduce antibiotic usage in finishing pigs by genetic selection. AB_mg_ and UD_Pen_ are weight-based methods to quantify antimicrobial usage, whereas TI_ADD_ accounts for the frequency of antibiotic administrations. Expressing antibiotic usage in mg active product (AB_mg_) or ml product (AB_ml_) might be interesting as description for the used antibiotics, however it seems less relevant for estimates of h^2^: 10 mg (or ml) of antibiotic x is not necessarily equivalent to 10 mg (or ml) of antibiotic y. Therefore, UD and/or TI_ADD_ seem better parameters for quantifying antibiotic usage, since they take into account the mg of active compound administered per kg of animal at risk.

Heritability estimates remained relatively consistent in the cross-validations (Table 4), implying that h^2^ estimates were robust against random sampling of data and also the reduction in data points.

Heritability estimates of N_treatment-all_ and N_treatment-respiratory_ were 25% and 3%, respectively. Putz et al. (21) found a heritability of 13-29% for the number of antibiotic treatments (depending on calculation method) at individual pig level, while Henryon et al. (20) found heritability estimates of 10-19% for time until first treatment on a logarithmic scale for different disease categories. Heritability estimates (at pen level) for mortality, FI and FCR and (at the individual level) for ADG and Meat% were similar to estimates reported in literature (summarised in Rothschild and Ruvinsky (34)).

Common environmental effects (c^2^) were moderate to high for traits related to antibiotic usage. This was expected since disease prevalence and antimicrobial usage are known to show seasonal variation in pigs, depend on the presence of infectious agents and are linked to the stable climate (1,35). Interestingly, c^2^-estimates were consistently higher in the respiratory problems dataset, meaning these environmental effects within a compartment are larger for respiratory problems. Indeed, the stable climate is well known to play a pivotal role for the respiratory health of pigs (36).

Predictive abilities using five-fold cross-validation were highest and significantly larger than zero for UD_Pen_, even when only distant genetic relationships (across-family strategy) were exploited. This indicates the potential of UD_Pen_ for use in pig breeding programs. Applying one generation of single-trait selection to reduce UD_Pen-all_ (additive genetic variance is 3.1 mg^2^ active compound/kg^2^ pig) with a selection intensity of 1, a generation interval of 3 years and an accuracy of 0.50 would yield an annual reduction of UD_Pen-_ all by 0.29 mg active compound/kg pig when using the breeder’s equation and assuming an equal potency between products.

Genetic correlations of UD_Pen_ and N_treatment_ with mortality at pen level were low to moderately high (0.08-0.60 ;Table 3) but consistently favourable and significantly larger than 0 for UD_Pen_ (Table 3). This genetic correlation indicates that when selecting pigs for a low (favourable) breeding value for antibiotic usage (based on UDPen), mortality at the pen level would simultaneously decrease as a correlated response. We believe this is a very interesting result, as mortality is one of the most important and ignored traits in pig breeding programs. Furthermore, no strong adverse genetic correlations were found with production traits like FI, ADG and FCR. However, we would like to note that not all genetic correlations were significantly different from zero. Only between N_treatment_ and Meat% unfavourable genetic correlations were found (0.45 and 0.11), whereas low favourable genetic correlations were found between UD_pen_ and ADG (−0.30 and -0.21) and FCR (0.25 and 0.26). These results indicate that UD_Pen_ or N_treatment_ can be included in current breeding programs without adversely affecting other economically important traits. Moreover, breeding pigs for lower UD_Pen_ or N_treatment_ will not only decrease antimicrobial usage by increasing general pig health, but may also decrease costs (purchase, labour) for pig farmers and reduce the risk of antibiotic resistance (11,15).

### Limitations

The present study is based on data from finishing pigs at pen level in two experimental farms considered representative for the Belgian pig sector. The dataset contained 2238 full-sib pens, which is a reasonable size to estimate heritabilities, but might still be insufficient to accurately estimate genetic correlations although standard errors of genetic correlations were low to moderate (Table 3). Furthermore, no data was available on antibiotic usage of these pigs during the suckling and/or nursery period, making it impossible to estimate genetic correlations between age categories. The results of this study are however very encouraging and can form the basis for further studies in different environments, using larger datasets from different age categories and/or using different breeds to confirm these results.

### Future perspectives

In the present study, data on antibiotic usage were recorded on paper. However, recent technology allows automated recording of treatments of individual pigs in real time using RFID tags. Such new precision livestock farming solutions may offer a user-friendly solution to record antibiotic usage in large scale pig breeding programs. Individual data may increase accuracy of breeding values and, hence, genetic progress. Moreover, individual recording is required in pens of non-siblings in order to exploit pedigree or genomic relationships.

Individual tracking using RFIDs would also allow to study the suckling and weaning periods during which most antibiotics are used. Sarrazin et al. (2) and Sjölund et al. (31) found that about 25% of all antibiotics is used in suckling pigs, 65% in weaners and only about 10% in fattening and/or breeding pigs. Although positive associations were found between antimicrobial administrations in the different age categories (2,31), the genetic correlations of antibiotic usage between age categories is still unknown. Recording antimicrobial usage of individual pigs throughout their lifespan seems valuable to get a full view on a pig’s genetic potential regarding antimicrobial usage. Moreover, such data would allow to estimate genetic correlations of antibiotic usage between different age categories.

## 5. Conclusions

This study shows that new phenotypes, derived from on-farm drug records at pen level from Piétrain sired crossbred finishing pigs in a commercial environment, were heritable. Moreover, low to moderate favourable genetic correlations were found with mortality and (small) favourable genetic correlations were estimated with production traits such as average daily gain and feed conversion ratio. Hence, these new phenotypes are promising traits to be included in the pig breeding programs, although more research is still necessary at this point. Further improvements are expected by recording antibiotic usage on an individual level throughout a pig’s lifespan via emerging precision livestock farming technologies.

## Supporting information

Table S1. Extra information on administered antibiotics

## Supplementary Materials

The following are available online at www.mdpi.com/xxx/s1, Figure S1: title, Table S1: title, Video S1: title.

## Author Contributions

WG analysed the data and wrote the manuscript. WG, DM, RM, JD, SJ and NB designed and conceived this study. DM, RM, JD, SJ and NB critically reviewed the analyses and the manuscript. All authors read and approved the final manuscript.

## Institutional Review Board Statement

Not applicable.

## Informed Consent Statement

Not applicable.

## Data Availability Statement

The dataset will be made accessible upon motivated request and is available upon request during the review process.

## Acknowledgments

The authors would like to acknowledge Dr. Bénédicte Callens for her help in constructing antibiotic parameters. Furthermore we would like to thank Stan Cardinaels, Pauline Debontridder, Joost Matthijssen, Ben Van Hemelen en Margot Verstreken for digitalizing antibiotic records.

## References

1. Lekagul A, Tangcharoensathien V, Yeung S. Patterns of antibiotic use in global pig production: A systematic review. Vol. 7, Veterinary and Animal Science. Elsevier B.V., 2019;100058.

2. Sarrazin S, Joosten P, Van Gompel L, Luiken REC, Mevius DJ, Wagenaar JA, et al. Quantitative and qualitative analysis of antimicrobial usage patterns in 180 selected farrow-to-finish pig farms from nine European countries based on single batch and purchase data. J Antimicrob Chemother. 2019;74(3):807–16.

3. Aarestrup F. Get pigs off antibiotics. Nature. 2012;486(7404):465–466.

4. European Union. Moving towards a more healthy and sustainable EU food system, a corner stone of the European Green Deal. 2020. Available from https://ec.europa.eu/info/strategy/priorities-2019-2024/european-green-deal/actions-being-taken-eu/farm-fork_en

5. Maron DF, Smith TJS, Nachman KE. Restrictions on antimicrobial use in food animal production: An international regulatory and economic survey. Global Health. 2013;9(1).

6. Postma M, Vanderhaeghen W, Sarrazin S, Maes D, Dewulf J. Reducing Antimicrobial Usage in Pig Production without Jeopardizing Production Parameters. Zoonoses Public Health. 2017;64(1):63–74.

7. European Medicines Agency. European Surveillance of Veterinary Antimicrobial Consumption (ESVAC) [Internet]. European Surveillance of Veterinary Antimicrobial Consumption (ESVAC). 2019 [cited 2020 Jun 2]. Available from: https://www.ema.europa.eu/en/veterinary-regulatory/overview/antimicrobial-resistance/european-surveillance-veterinary-antimicrobial-consumption-esvac

8. FAGG. SANITEL-MED [Internet]. 2020. Available from: https://www.fagg-afmps.be/nl/SANITEL-MED

9. Maes D, Dewulf J, Boyen F, Haesebrouck F. Disease identification and management on the pig farm. In 2018; 77–99.

10. Nakov D, Hristov S, Stankovic B, Pol F, Dimitrov I, Ilieski V, et al. Methodologies for assessing disease tolerance in pigs. Vol. 5, Frontiers in Veterinary Science. Frontiers Media S.A., 2019;329

11. Guy SZY, Li L, Thomson PC, Hermesch S. Genetic parameters for health of the growing pig using medication records. Proc 11th World Congr Genet Appl to Livest Prod. 2018;

12. Vissche AH, Janss LLG, Niewold TA, de Greef KH. Disease incidence and immunological traits for the selection ofhealthy pigs a review. Vol. 24, Veterinary Quarterly. 2002;29–34.

13. Gross WG, Siegel PB, Hall RW, Domermuth CH, DuBoise RT. Production and persistence of antibodies in chickens to sheep erythrocytes. 2. Resistance to infectious diseases. Poult Sci. 1980;59(2):205–10.

14. Nejsum P, Roepstorff A, Jørgensen CB, Fredholm M, Göring HHH, Anderson TJC, et al. High heritability for Ascaris and Trichuris infection levels in pigs. Heredity (Edinb). 2009;102(4):357–64.

15. Putz A. Quantifying resilience in sows and wean-to-finish pigs. Graduate Theses and Dissertations. Iowa State University; 2019.

16. Flori L, Gao Y, Laloë D, Lemonnier G, Leplat JJ, Teillaud A, et al. Immunity traits in pigs: Substantial genetic variation and limited covariation. PLoS One. 2011;6(7).

17. Clapperton M, Diack AB, Matika O, Glass EJ, Gladney CD, Mellencamp MA, et al. Traits associated with innate and adaptive immunity in pigs: Heritability and associations with performance under different health status conditions. Genet Sel Evol. 2009;41(1).

18. Bishop SC, Woolliams JA. Genomics and disease resistance studies in livestock. Livest Sci. 2014;166(1):190–8.

19. Guy S, Thomson PC, Hermesch S. Selection of pigs for improved coping with health and environmental challenges: Breeding for resistance or tolerance? Vol. 3, Frontiers in Genetics. 2012;281

20. Henryon M, Berg P, Jensen J, Andersen S. Genetic variation for resistance to clinical and subclinical diseases exists in growing pigs. Vol. 73, Animal Science. 2001; 375–387

21. Putz AM, Harding JCS, Dyck MK, Fortin F, Plastow GS, Dekkers JCM. Novel resilience phenotypes using feed intake data from a natural disease challenge model in wean-to-finish pigs. Front Genet. 2019;660.

22. Timmerman T, Dewulf J, Catry B, Feyen B, Opsomer G, Kruif A de, et al. Quantification and evaluation of antimicrobial drug use in group treatments for fattening pigs in Belgium. Prev Vet Med. 2006;74(4):251–63.

23. Jensen VF, Jacobsen E, Bager F. Veterinary antimicrobial-usage statistics based on standardized measures of dosage. Prev Vet Med. 2004; 64(2–4):201–15.

24. Callens B, Persoons D, Maes D, Laanen M, Postma M, Boyen F, et al. Prophylactic and metaphylactic antimicrobial use in Belgian fattening pig herds. Prev Vet Med. 2012;106(1):53–62.

25. AMCRA. Antibioticadoseringenlijst varkens maart 2020 [Internet]. 2020. Available from: https://www.amcra.be/swfiles/files/AB-doseringenlijst_varkens_maa-2020_406.pdf

26. Misztal I, Tsuruta S, Lourenco D, Aguilar I, Legarra A, Vitezica Z. Manual for BLUPF90 family of programs. Athens Univ Georg. 2014;

27. Muñoz F, Sanchez L. breedR: Statistical Methods for Forest Genetic Resources Analysts. 2019.

28. Legarra A, Robert-Granié C, Manfredi E, Elsen JM. Performance of genomic selection in mice. Genetics. 2008;180(1):611–8.

29. Lekagul A, Tangcharoensathien V, Yeung S. The use of antimicrobials in global pig production: A systematic review of methods for quantification. Vol. 160, Preventive Veterinary Medicine. Elsevier B.V., 2018; 85–98.

30. Trauffler M, Griesbacher A, Fuchs K, Köfer J. Antimicrobial drug use in Austrian pig farms: Plausibility check of electronic on-farm records and estimation of consumption. Vet Rec. 2014;402

31. Sjölund M, Postma M, Collineau L, Lösken S, Backhans A, Belloc C, et al. Quantitative and qualitative antimicrobial usage patterns in farrow-to-finish pig herds in Belgium, France, Germany and Sweden. Prev Vet Med. 2016;41–50

32. Abdelsayed M, Haile-Mariam M, Pryce JE. Genetic parameters for health traits using data collected from genomic information nucleus herds. J Dairy Sci. 2017;100(12):9643–55.

33. Gunia M, David I, Hurtaud J, Maupin M, Gilbert H, Garreau H. Resistance to infectious diseases is a heritable trait in rabbits. J Anim Sci. 2015;93(12):5631–8.

34. Rothschild, M. F., and A. Ruvinsky, eds. The genetics of the pig. CABI, 2011.

35. Chmielowiec-Korzeniowska A, Tymczyna L, Babicz M. Assessment of selected parameters of biochemistry, hematology, immunology and production of pigs fattened in different seasons. Arch Anim Breed. 2012;55(5):469–79.

36. Zimmerman JJ, Karriker LA, Ramirez A, Schwartz KJ, Stevenson GW, Zhang J. Diseases of Swine. Wiley-Blackwell, 2019.

