## Supplementary material for "High heritabilities for antibiotic usage show potential to breed for disease resistance in finishing pigs": Table S1. Extra information on administered antibiotics

Additional File 1 Table S1. Extra information on administered antibiotics. The LA-factor (Long Acting) is used to calculate treatment incidences to account for a possible long acting effect (AMCRA, 2020). Only parenterally administered antibiotics were used in this study.

| Class antibiotics | Active compound | Commercial drug name | Concentration (mg/ml) | LA-factor |
| --- | --- | --- | --- | --- |
| Amfenicoles | Florfenicol | Florfenikel®, Kela | 300 | 2 |
| Aminopenicillines | Amoxicillin | Vetrimoxin Long Acting®, Ceva Santé Animale NV | 150 | 1 |
| Aminosides | Paromomycin | Gabbrovet®, Ceva Santé Animale NV | 175 | 1 |
| Cephalosporins | Ceftiofur | Ceftiosan®, Alfasan International BV | 50 | 1 |
| Cephalosporins | Cefquinome | Cobactan 2,5%®, Intervet International BV | 25 | 1 |
| Cephalosporins | Ceftiofur | Excenel Flow®, Zoetis Belgium SA | 50 | 1 |
| Fluoroquinolones | Enrofloxacin | Fenoflox®, Chanelle Pharmaceuticals Manufacturing Ltd | 50 | 1 |
| Lincosamides | Lincomycin | Lincomycine-VMD®, VMD NV | 100 | 1 |
| Macrolides | Tulathromycin | Draxxin®, Zoetis Belgium SA | 100 | 9 |
| sulfonamides-trimethoprim | Trimethoprim  Sulfadoxine | Dofatrim-ject®, Dopharma Research BV | 40  200 | 1 |
